# Sex differences in brain structure: An autism twin study on restricted and repetitive behaviors

**DOI:** 10.1101/334367

**Authors:** Annelies van’t Westeinde, Élodie Cauvet, Roberto Toro, Ralf Kuja-Halkola, Janina Neufeld, Katell Mevel, Sven Bölte

## Abstract

Females with autism spectrum disorder have been reported to exhibit fewer and less severe restricted and repetitive behaviors and interests compared to males. This difference might indicate sex specific alterations of brain networks involved in autism symptom domains, especially within cortico-striatal and sensory integration networks. This study used a well-controlled twin design to examine sex differences in brain anatomy in relation to repetitive behaviors. In 75 twin pairs (n=150, 62 females, 88 males) enriched for autism spectrum disorder (n=32), and other neurodevelopmental disorders (n =32), we explored the association of restricted and repetitive behaviors and interests – operationalized by the Autism Diagnostic Interview-Revised (C domain) and the Social Responsiveness Scale-2 (Restricted Interests and Repetitive Behavior subscale), with cortical volume, surface area and thickness of neocortical, sub-cortical and cerebellar networks. Cotwin control analyses revealed within-pair associations between RRBI symptoms and the right intraparietal sulcus and right orbital gyrus in females only. These findings endorse the importance of investigating sex differences in the neurobiology of autism symptoms, and indicate different etiological pathways underlying restricted and repetitive behaviors and interests in females and males.

## Introduction

Autism spectrum disorder (ASD) is a neurodevelopmental condition of complex origin, defined by challenges in social communication and interaction alongside restricted and repetitive behaviors and interests (RRBIs) that cause significant impairment in daily life functioning (Hallmayer 2011; Sandin et al. 2014; Colvert et al. 2015; de Schipper et al. 2015). A markedly skewed sex distribution has been consistently reported in ASD, despite recently improved recognition of autism in females (Jensen et al. 2014); the ratio is still estimated around 2-3 (males):1 (females) (Jensen et al. 2014; Baxter et al. 2015; Idring et al. 2015). The sex bias in ASD is hypothesized to arise from a female protective effect alongside male risk factors (Lai et al. 2015). In addition, differences might exist in the underlying etiology and symptom presentation of ASD in females, which could be associated with both reduced risk of developing ASD, as well as failure of recognizing ASD in females (Lai et al. 2017). Thus, investigating sex differences in the neurobiology associated with ASD symptom domains is crucial to understand pathways leading to ASD in both men and women. Moreover, recent Research Domain Criteria (RDoC) guidelines by the National Institutes of Health advise to quantify symptoms and functional domains for research purposes, rather than condensing them as categorical entities, in order to effectively investigate full variation of behaviors from typical to atypical. The latter is particularly relevant for ASD, since autistic traits have been found to be continuously distributed in the general population (Ronald et al. 2006; Robinson et al. 2016).

Sex differences in symptom presentation of ASD have predominantly been reported in the domain of RRBIs. Despite some inconsistencies (Carter et al. 2007; Holtmann et al. 2007), most studies found reduced frequency and severity of RRBIs in females (Bolte et al. 2011; Mandy et al. 2012; Van Wijngaarden-Cremers et al. 2014), in particular less special, narrow, and intense interests (Frazier et al. 2014). These differences might be caused by divergent etiological pathways of RBBIs in autism, including underlying brain anatomy. However, so far, the brain anatomy associated with RRBIs has mainly been studied in ASD males. RRBIs have been associated with cortico-striatal circuits that connect lateral orbitofrontal, anterior cingulate cortex and precentral motor regions to the striatum (Langen et al. 2011). In autistic males, the majority of neuroanatomical studies of RRBIs focused on subcortical areas. Here, the most conclusive finding was regional enlargement in both children and adults, in particular in the striatum, namely the caudate nucleus (Sears et al. 1999; Hollander et al. 2005; Haznedar et al. 2006; Rojas et al. 2006; Langen et al. 2007) and the globus pallidus (Schuetze et al. 2016). In addition, volume enlargements were found for the thalamus (Lin et al. 2015), amygdala and precentral gyrus (Rojas et al. 2006). However, some reductions in volumes were observed as well, for example in the inferior frontal gyri and cerebellum (Rojas et al. 2006).

So far, only one study has addressed sex differences in brain anatomy related to RRBIs, on the ABIDE dataset assessing 25 females and 25 males with ASD. The authors reported that gray matter of motor regions could discriminate boys from girls with ASD (Supekar and Menon 2015). In addition, only in girls RRBIs were related to increased gray matter of the motor cortex, the supplementary motor area and Crus 1 subdivision of the cerebellum, while correlating with the right putamen in boys (Supekar and Menon 2015). These findings indicate a different relationship between brain anatomy and RRBIs for males and females with ASD, thus potentially pointing to divergent etiological pathways to inflexible behaviors between the sexes.

More generally, ASD is associated with environmental, shared and non-shared, as well as genetic components which likely contribute to the heterogeneity in the etiology (Hallmayer 2011; Sandin et al. 2014; Colvert et al. 2015). Using a co-twin control design, enables to study gene-independent neuroanatomical variation associated with RRBI symptoms, for example life-long presence of RRBI symptoms themselves might alter brain-structure, and thus comprise a non-shared environment factor between the twins. In addition, a co-twin design reduces heterogeneity caused by age, gender, socio-economic background, other shared-environment and genetic make-up. Previous twin studies have observed structural changes in brain regions relevant for RRBIs, including the caudate nucleus, pre- and postcentral gyri (Kates et al. 1998) and cerebellum (Kates et al. 1998, 2004; Mitchell et al. 2009) (see (Mevel et al. 2015) for a review). However, none of these studies has addressed sex differences or RRBI symptoms directly.

As part of the Roots of Autism and ADHD Twin Study Sweden (RATSS; (Bölte et al. 2014)), the objective of this explorative study was to examine sex differences in the neuroanatomy of regions of interest in relation to a dimensional estimate of RRBIs using a within-pair twin design. Surface-based estimates, including volume, surface area and thickness of regions relevant for RRBIs were analyzed in same-sex twins aged 9 to 23 years. This clinically enriched sample consisted of typically developing twin pairs, in addition to pairs being concordant or discordant for ASD and other neurodevelopmental conditions.

## Materials and methods

### Participants

Complete twin sample characteristics are presented in Table 1. Informed written consent was obtained from all participants and/or their legal guardians according to the Declaration of Helsinki. The RATSS project and the current study are approved by the regional Ethical Review Board. Of in total n=288 twins included in RATSS to date, N=261 had completed MRI scanning, from which we only included same-sex pairs for which high quality image scans for both twins were available. These inclusion criteria resulted in a sample consisting of 75 same-sex pairs (n=150, age: 9 – 23 years), of which 44 were male pairs (mean age 15.9 years) and 31 female pairs (mean age 16.4 years), and 46 monozygotic and 29 dizygotic pairs. Zygosity was determined with DNA testing for most pairs, apart from 5 male and 2 female pairs who’s zygosity was established with a questionnaire. The sample included 32 twins with ASD (20 males, 12 females) from 20 ASD discordant (only one twin of a pair received an ASD diagnosis, the other not) and 6 ASD concordant pairs (both twins of a pair received ASD diagnosis), 34 twins with ADHD (23 males, 11 females), 25 twins with other neurodevelopmental disorders (16 males, nine females) and 70 without a diagnosis (40 males, 30 females).

**Table 1.** Whole twin sample and sex specific characteristics.

| **Demographics** | **All**<br>**(N=150)** | **Males**<br>**(N=88)** | **Females**<br>**(N=62)** | **P** |
| --- | --- | --- | --- | --- |
| Number of Pairs | 75 | 44 | 31 |  |
| Age mean (*SD*) | 16.11 (3.31) | 15.90 (3.17) | 16.39 (3.51) | .44 |
| Age range | 9 - 23.69 | 10.69 - 23.34 | 9 - 23.69 |  |
| Zygosity MZ/DZ | 92 / 58 | 50 / 38 | 42 / 20 | .20 |
| ASD diagnosis | 32 | 20 | 12 | .77 |
| ASD diagnosis per MZ / DZ | 15 / 17 | 7 / 13 | 8 / 4 |  |
| Pairs concordant for ASD | 6 | 3 | 3 |  |
| Pairs discordant for ASD | 20 | 14 | 6 |  |
| Pairs ASD concordant per MZ/DZ | 4 / 2 | 1 / 2 | 3 / 0 |  |
| Pairs ASD discordant per MZ/DZ | 7 / 13 | 5 / 9 | 2 / 4 |  |
| ADHD diagnoses | 34 | 23 | 11 | 0.24 |
| Other neurodevelopmental disorders | 25 | 16 | 9 | 0.66 |
| Psychiatric diagnoses | 24 | 14 | 10 | 1 |
| No diagnosis | 70 | 40 | 30 | 0.74 |
| Age ASD discordant pairs | 15.98 (3.50) | 17.70 (3.16) | 15.19 (3.41) | 0.033* |
| range | 10.69-23.34 | 12.86-21.52 | 10.69-23.34 |  |
| Age ASD concordant pairs | 16.28 (4.08) | 19.30 (3.47) | 13.26(1.65) | 0.004** |
| range | 11.09-23.69 | 16.34-23.69 | 11.09-14.43 |  |
| Age non-ASD pairs | 15.91(3.06) | 15.33(3.14) | 16.36(2.94) | 0.093 |
| range | 9.00-22.77 | 9.00-22.17 | 10.95-22.77 |  |

*MZ= monozygotic, DZ= dizygotic. Concordant: both twins with ASD diagnosis, Discordant: only one sibling with ASD diagnosis. P-values from chi-square test comparing males and females.*

### Measures

#### Behavioral assessments

The comprehensive phenotypic assessment protocol of RATSS is described elsewhere in detail (Bölte *et al*., 2014). Briefly, clinical consensus diagnosis of ASD and other neurodevelopmental disorders or absence of clinical diagnosis was based on DSM-5 criteria (Association AP. 2013) by three experienced clinicians, supported by information from the Autism Diagnostic Interview - Revised (ADI-R; (Rutter, M. et al. 2003), the Autism Diagnostic Observation Schedule-2 (Lord, C. et al. 2012), the Kiddie Schedule for Affective Disorders and Schizophrenia (Kaufman et al. 1997) or the Diagnostic Interview for ADHD in Adults (Kooij, J.J.S. 2010). Full-scale IQ was assessed based on performance on the Wechsler Intelligence Scales for Children and Adults, Fourth Editions (Wechsler, D. 2003; Wechsler, D. et al. 2008). Handedness was assessed using the Edinburgh Handedness Inventory (Oldfield 1971), which estimates laterality in everyday life on a scale from - 100% (left handed) to +100% (right handed).

For our main analyses, the frequency and severity of RRBIs was determined by the RRBI subscale (domain C) of the ADI-R, using item codes for lifetime symptom presentation (“ever”). In the diagnostic algorithm of the ADI-R, the RRBI subscale comprises 8 items scored 0 to 2, with “0” indicating no RRBIs typical of autism, “1” RRBIs typical of autism, but mild, or “2” RRBIs prototypic of autism) [max. total score = 16]. The diagnostic cut-off for presence of clinically relevant RRBIs indicating ASD on the total score ≥ 2 (n=41 in our sample). On the ADI-R, scoring even 1 point has clinical value. 37 pairs were discordant for RRBI symptoms; these pairs had a within-pair difference on RRBIs of at least 1 point. Please, see Table 2 and Supplementary fig. 1A, for the ADI-R RRBI score distribution in our twin sample. Further, post-hoc analyses addressed the robustness in terms of operationalization and time frame using a different RRBI estimate, the Restricted Interests and Repetitive Behavior (RRB) subscale of the Social Responsiveness Scale-2 (SRS-2) standard child or adult version (Constantino 2005). The SRS-2 assesses autistic-like behaviors and quantifies its severity focusing on the past six-month, as opposed to the life-time symptom assessment of the ADI-R. Raw scores on the SRS-2-subscale autism mannerisms were retrieved, that comprises of 12 items scored 0 to 3 on a Likert-scale [max. total score = 36], with higher scores indicating the presence of more autistic mannerisms, including repetitive behaviors and restricted interests. In addition, in order to test the specificity of potential brain anatomical findings to RRBI’s, against social communication & cognition aspects of autism, we also used the social cognition subscale from the SRS-2, comprising 12 items [max. total score = 36] assessing past 6 months social cognition abilities, as well as the reciprocal interaction domain (domain A) from the ADI-R, comprising 16 items assessing life-time reciprocal interactions [max. total score = 32]. For all subscales, a higher score indicates more problems with RRBI’s, social cognition and reciprocal interaction respectively.

**Table 2.** Twin sample characteristics for the behavioral variables.

| **A) Mean Scores** | **All**<br>**(N=150)** | **Males**<br>**(N=88)** | **Females**<br>**(N=62)** | **P** |
| --- | --- | --- | --- | --- |
| IQ mean (*SD*) | 97.86 (15.65) | 97.22 (14.42) | 98.77 (17.31) | .71 |
| range | 62-142 | 62-123 | 63-142 |  |
| ADI-R RRBI mean (*SD*) | 1.02 (1.62) | 1.10 (1.81) | 0.90 (1.33) | .77 |
| range | 0-10 | 0-10 | 0-5 |  |
| ADI-R social interaction mean (*SD*) | 4.08 (5.13) | 4 (5.41) | 4.19 (4.76) | 0.60 |
| range | 0-25 | 0-25 | 0-16 |  |
| Pairs ADI-R RRBI discordant* | 37 | 21 | 16 |  |
| SRS-2 total score mean (*SD*) | 41.66 (30.97) | 40.65 (29.50) | 43.10 (33.15) | .93 |
| range | 0-131 | 4-131 | 0-130 |  |
| SRS-2 RRB mean (*SD*) | 5.85(6.25) | 6.08 (6.41) | 5.52 (6.07) | 0.60 |
| range | 0-26 | 0-26 | 0-23 |  |
| SRS-2 social cognition mean (*SD*) | 7.66 (6.50) | 7.20(5.88) | 8.31(7.29) | 0.59 |
| range | 0-29 | 0-23 | 0-29 |  |
| Handedness (mean) | 66.07 | 66.72 | 65.15 | .20 |
| **B) Within-pair differences** |  |  |  |  |
| IQ mean (*SD*) | 10.15 (8.81) | 9.84 (9.48) | 10.58 (7.90) | .38 |
| range | 0-39 | 0-39 | 0-31 |  |
| ADI-R RRBI mean (*SD*) | 1.05 (1.36) | 1.022 (1.36) | 1.097 (1.40) | .79 |
| range | 0-5 | 0-5 | 0-5 |  |
| ADI-R social interaction mean (*SD*) | 2.93(4.27) | 3.23(5.27) | 2.52(3.85) | 0.89 |
| range | 0-25 | 0-25 | 0-15 |  |
| SRS-2 total score mean (*SD*) | 23.05 (26.22) | 23.39 (25.55) | 22.58 (27.57) | .99 |
| range | 0-101 | 0-96 | 0-101 |  |
| SRS-2 RRB mean (*SD*) | 4.92 (5.77) | 5.07 (5.93) | 4.71 (5.62) | .83 |
| range | 0-22 | 0-22 | 0-22 |  |
| SRS-2 SCOG mean (*SD*) | 5.4(5.35) | 5.32(4.79) | 5.52(6.14) | 0.71 |
| range | 0-22 | 0-19 | 0-22 |  |

*\*ADI-R C discordancy refers to the number of pairs that have a within-pair difference on the ADIR C score of > 0 and therefore contribute most to the within-pair analyses.*
*SD: standard deviation. IQ: overall intellectual quotient measured with WISC/WAIS. Handedness: assessed with Edinburgh handedness inventory, a scale from -100/+100, where -100 represents totally left handed, and +100 totally right handed. P-values are given from chi-square tests comparing males with females.*

### Structural MRI

#### Image acquisition

T1-weighted images were acquired on a 3 Tesla MR750 GE scanner at the Karolinska Institutet MR center (Inversion Recovery Fast Spoiled Gradient Echo - IR-FSPGR, 3D-volume, 172 sagittal slices, 256×256, FOV 24, voxel size 1mm^3^, flip angle 12, TR/TE 8200/3.2, using a 32-channel coil array). T1-weighted acquisition was part of a 50-minute scanning protocol, preceded by a 5 to 7 minutes mock scan training for self-control of head movements. During the mock scan training participants were equipped with a sensor on their forehead measuring head-movement while they watched a cartoon movie on the screen via a mirror. With excess head movement (1.5 mm in any direction), the video would stop, thus providing feedback to the participant about their movement. Head movement reduced during the practice for most participants.

#### Surface-based neocortical and subcortical analyses: cortical volumetry, cortical thickness and surface area (Freesurfer 6)

Raw images were processed in Freesurfer 6 (http://surfer.nmr.mgh.harvard.edu/). The well-established standard pipeline was run on the original T1-weighted images (Dale et al. 1999; Fischl et al. 1999). Briefly, the intensity of the images was normalized, the brain was skull stripped and brain tissues were segmented. A white matter volume was generated, from which a surface tessellation was created. Meshes were constructed for gray and white matter out of approximately 150,000 vertices per hemisphere, then parcellated according to the Destrieux Atlas (Destrieux et al. 2010). Next, mean cortical thickness, volume and surface areas were obtained for each region in each hemisphere. Whole brain volume from FreeSurfer was used as a covariate in all surface- and volume-based analyses except for cortical thickness, because cortical thickness is less related to brain volume (Toro et al. 2008). After a quality control of the brain data processed from an initial 261 subjects that had completed MR scanning, 150 participants with three outputs each (cortical volume, surface area and cortical thickness) were retained in the final surface-based analyses. Quality control was done by visual inspection of the T1 images for presence of movement errors, accuracy of skull stripping, and accuracy of the FreeSurfer segmentation, i.e. inspecting if the pial and white matter surfaces accurately followed the intersection between brain/CSF and gray matter/white matter respectively. Minor segmentation errors, such as at the temporal poles, were tolerated, in particular with regard to the young age of the subject group. Subjects were given a score on movement and image quality, of 1 (no errors) – 4 (very severe movement), and only subjects with a score of 1 or 2 were included. Within pairs, RRBI’s (ADI-R) did not predict data quality (B=0.009, p=0.814) or movement scores (B=-0.009, p=0.889).

#### Volume-based cerebellar analysis: Gray and white matter regional volume (FSL)

Volumes of cerebellar white and gray matter were retrieved using volume-based morphometry. The 261 raw brain volumes were intensity normalized and the brain was extracted using AFNI’s 3dskullstrip. Skull stripped 3D images were segmented in 3 tissue types (Gray Matter-GM, White Matter-WM, Cerebral Spinal Fluid-CSF) using FAST (FMRIB’s Automated Segmentation Tool within FMRIB’s Software Library) which also corrects for spatial intensity variation. Segmented images were warped to MNI space using non-linear registration FNIRT from FSL. GM and WM volumes for the somato-motor cerebellar region were extracted from the intersection between the somato-motor regions in Buckner’s 7-network functional atlas (Buckner et al. 2011) and segmented individual volumes using a custom script in C. The same 150 individuals that had passed the surface-based quality control were included in the volume-based analyses. These 150 scans all had good segmentation quality in FSL, because the inclusion decision was based on the surface-based segmentation, which was more sensitive to quality of raw images.

#### ROI selection for the neocortical, subcortical and cerebellar RRBI neworks

RRBIs are presumed to rely on a wide network of regions involved in motor function and cognitive control of neocortical and subcortical areas, in particular cortico-striatal circuits (Langen et al. 2009). In the current study, we therefore focus on these cortico-striatal loops, motor regions and sensory integration areas, that have previously been associated with ASD, including pre- and postcentral motor regions, the striatum (Langen et al. 2011), the amygdala (Rojas et al. 2006) and sensory-motor integration areas in the posterior parietal cortex (Shafritz et al. 2008; Mostofsky and Ewen 2011), and areas involved in executive functioning in prefrontal areas (Langen et al. 2011) and cerebellum (Becker and Stoodley 2013). Based on these previous findings we selected a priori corresponding neocortical and subcortical regions of interest within the Destrieux atlas from Freesurfer (Destrieux et al. 2010). We included volumes, surface area and thickness of 16 regions that include the areas described above, namely the anterior cingulate cortex (ACC), lateral orbital sulcus, orbital gyrus, inferior frontal orbital gyrus, bilateral postcentral gyrus, postcentral sulcus, precentral gyrus, precentral inferior sulcus, precentral superior sulcus, central sulcus, superior frontal sulcus, superior frontal gyrus, supramarginal gyrus, superior parietal lobule, intra parietal sulcus and angular gyrus, as well as volumes of 5 subcortical regions, namely the bilateral caudate nucleus, globus pallidus, putamen, thalamus, amygdala, in addition to volume of the cerebellar cortex and white matter. We also included the volume of the somato-motor region of the cerebellum based on a functional connectivity atlas from FSL (Buckner et al. 2011).

### Statistical Analysis

All statistical analyses were performed in R (https://www.r-project.org/). All p-values are FDR corrected for Type I errors, significance threshold was set to p<.05. Sample size was comparable to recently published twin studies using similar co-twin designs reporting medium to large effect size (Wilson et al. 2015; Picchioni et al. 2017).

#### Sex differences in demographics

We first examined possibly confounding demographic differences between females and males. Statistical comparisons between the sexes were conducted using χ^2^ tests for categorical variables (zygosity, diagnosis) and Kruskal-Wallis tests for continuous variables (age, RRBIs, IQ, handedness scores). These tests yielded no between group significant differences (see Table 1).

#### Sex differences for neuroanatomy of the motor network in relation to RRBIs

The main analyses focused on within-pair differences in RRBIs as assessed with the ADI-R C domain, while post-hoc control analyses 1) cross-validated the findings with the RRB subscale from SRS-2 and 2) tested the specificity of the findings to RRBIs by controlling for social cognition and communication. Total brain volume was adjusted for when assessing cortical volume and surface area, but not thickness, and IQ was adjusted for in all models.

#### Twin/co-twin: Within-pair analyses

For the main analyses, a twin/co-twin design was implemented to investigate the association between RRBI’s on a dimensional scale (predictor) and the anatomy (outcome) of the regions of interest while controlling for unmeasured confounding factors shared within twin pairs (e.g. genetic factors, demographics etc.). MZ and DZ twins were collapsed in order to increase statistical power. Within-twin pair associations were estimated using a conditional linear regression model within the generalized estimating equations (GEE) framework, using the drgee package from R (Zetterqvist and Sjölander 2015). Herein, the difference in the exposure variable within a pair is correlated to the difference in the outcome variable within the same pair. A within-pair association was estimated by correlating the difference in the exposure variable to the difference in the outcome variable within a pair (see Supplementary fig. 2 and 3 for some examples). This within-pair relationship is calculated for all pairs resulting in an estimate for the average within-pair association between RRBIs and brain anatomy in the group. Thus, the association between RRBI and brain in our study was estimated by making use of dimensional discordancy within-twin pairs, i.e. differences within pairs on total scored points of RRBIs.

#### Step-wise within-pair analyses

Within-pair analyses were run in three sub-steps. First, the association between RRBIs and brain structure was assessed for males and females separately. Because the within-pair model assesses the associations within a pair, which are all same-sex, it was not possible to directly assess the interaction between sex and RRBIs. To compare the association between symptoms and structure in males and females, we computed Wald χ^2^ tests for each ROI that was associated with RRBIs in either males or females. A significant difference on a Wald-test indicates that the estimate of the association was different for the sexes.

#### Post-hoc analyses

Further, to test the robustness of the observed effects, models otherwise identical to the models of the main analyses were run with a different estimate of RRBIs, RRB subscale of the SRS-2, which addresses current as opposed to life-time symptoms. Finally, specificity of the results toward RRBIs were tested by adding different autism symptom domains as covariates in the model, including the social cognition subscale from SRS-2, and reciprocal social interaction domain from the ADI-R, to control for highly correlated symptoms that might have confounded the observed effects. Additional analyses were run to control for interaction effects between age and RRBIs on brain anatomy showing significant associations for the right postcentral gyrus, superior precentral sulcus and superior parietal sulcus, i.e. areas which were not associated with RRBIs in our study (Supplementary Tables).

## Results

### 1. Sex differences in RRBIs and autistic traits

Males and females did not differ on overall RRBI symptom severity, other autistic symptoms and traits and IQ. Further, no within pair differences between the sexes were observed for any of these variables (Table 2).

### 2. Twin/co-twin: Within-pair differences in RRBIs associated to within-pair differences in neuroanatomy of the motor network

#### 2.1 Main within pair effects of RRBIs (ADI-R) on brain anatomy for males and females

Main results are presented in Table 3A & B. When splitting the sample by sex and controlling for IQ, within-pair increases in RRBI symptoms were related to increased thickness of the right intraparietal sulcus in females only (see Figure 1 and Supplementary Figure 3). No other significant associations were observed. However, there was a trend for reduced surface area in the same region. Moreover, there were associations at trend level that followed a similar pattern in females, namely increased thickness of the right orbital gyrus, and right inferior frontal orbital gyrus, and reduced surface area of the left superior frontal gyrus. There was also a trend for increased surface area of the right middle frontal gyrus. In males, on the other hand, no significant within-pair associations between RRBIs and brain anatomy were observed.

**Figure 1:**
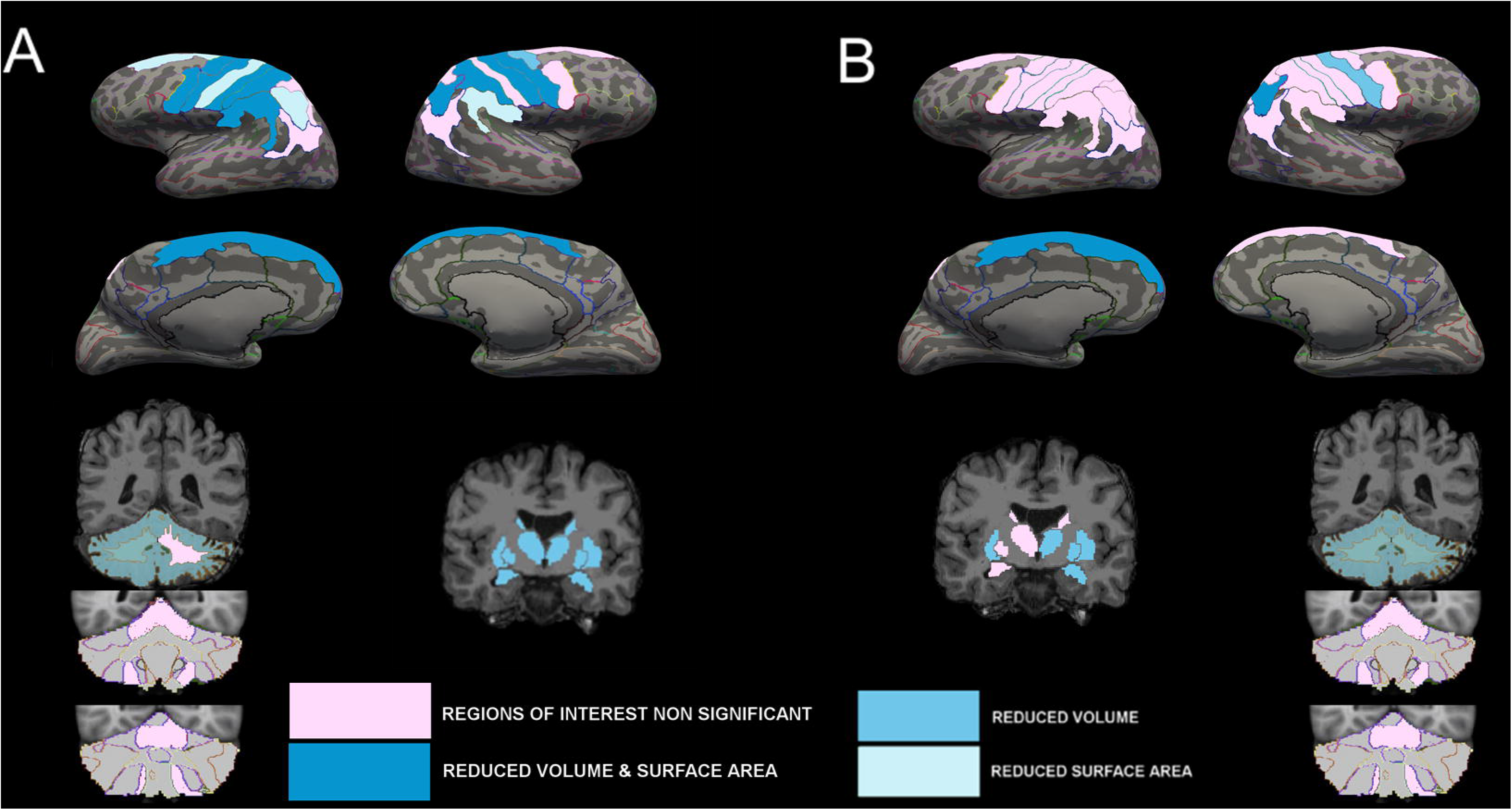
Brain region associated with restricted and repetitive behaviors and interests in females. Within-pair association between ADI-R C and brain structure. The area that was significantly associated with RRBIs is displayed blue: increased thickness of the right intraparietal sulcus in females. Areas not significantly associated with RRBIs, but included in our regions of interest, are displayed in pink.

**Table 3A.**
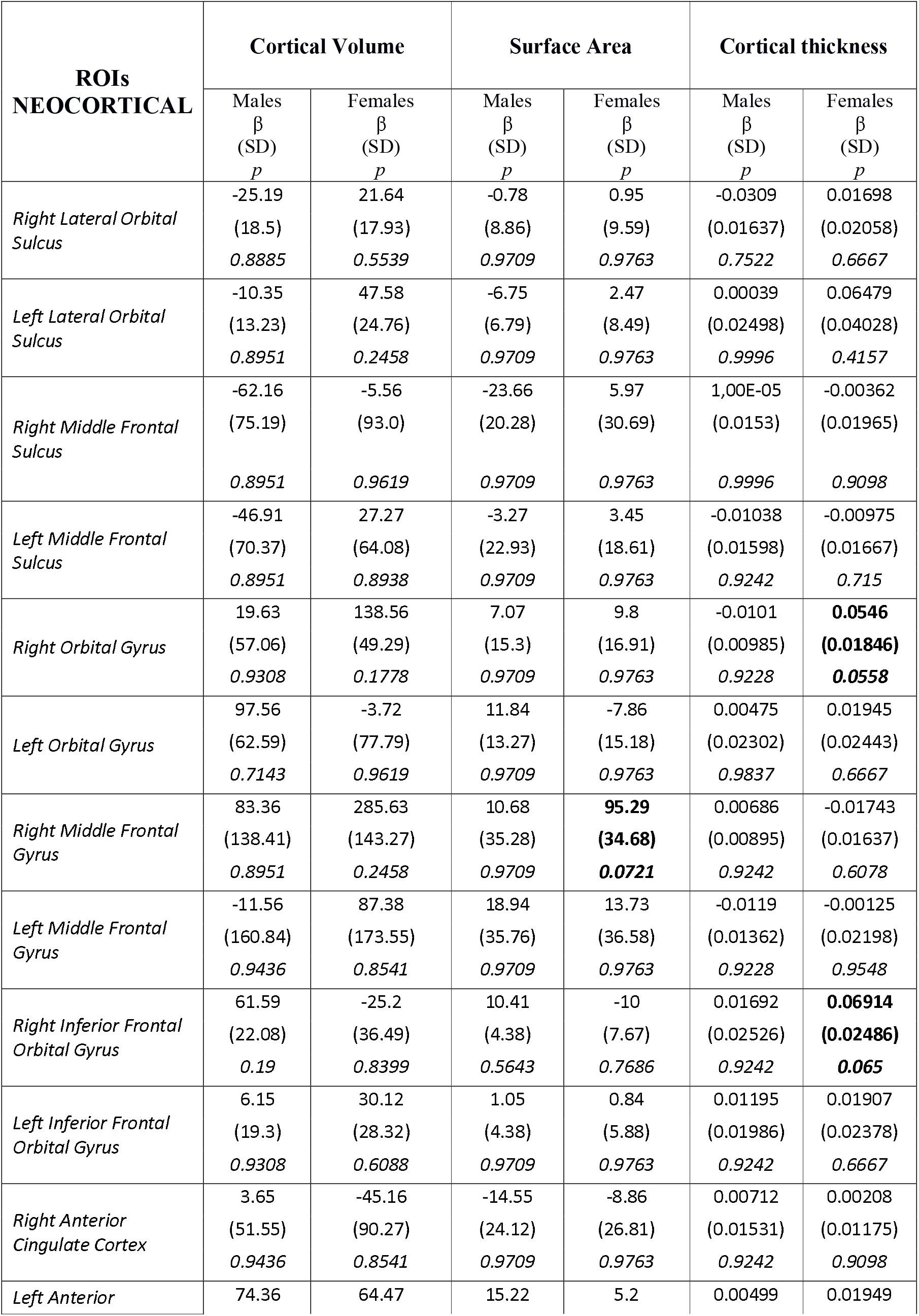
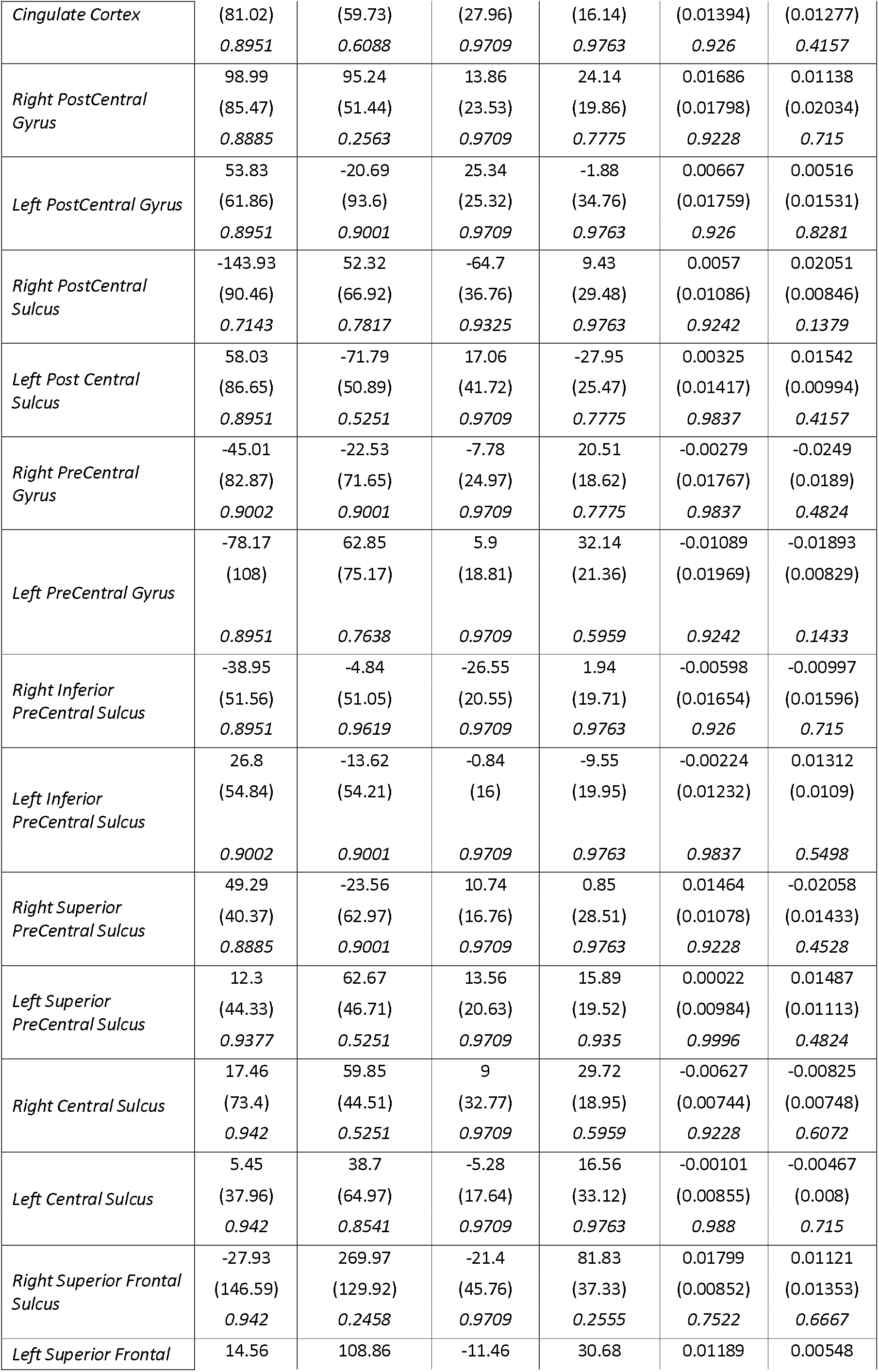
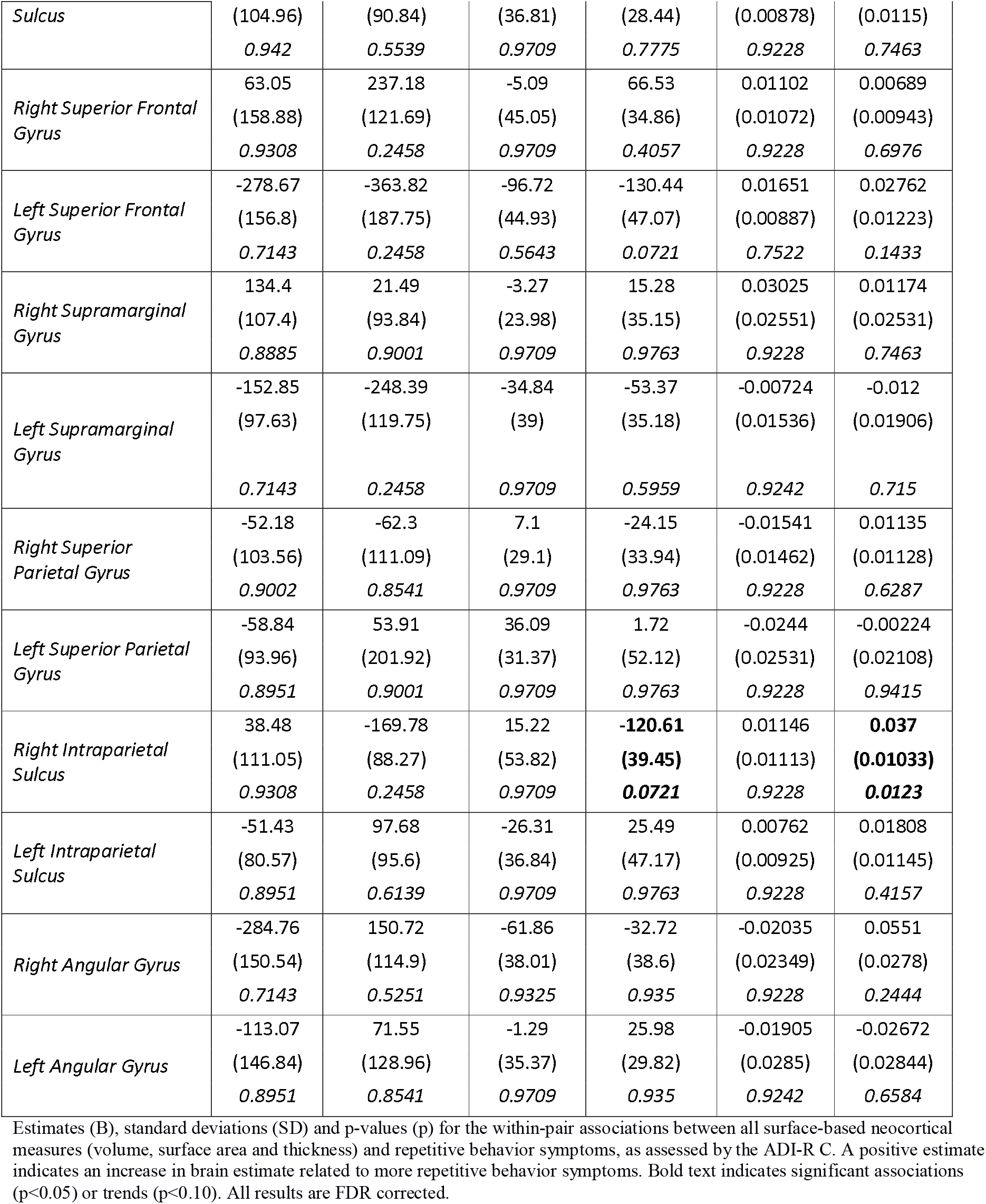
Twin model associations between cortical volume, surface area and thickness of neocortical regions of interest (ROIs) and RRBI symptoms.

**Table 3B.**
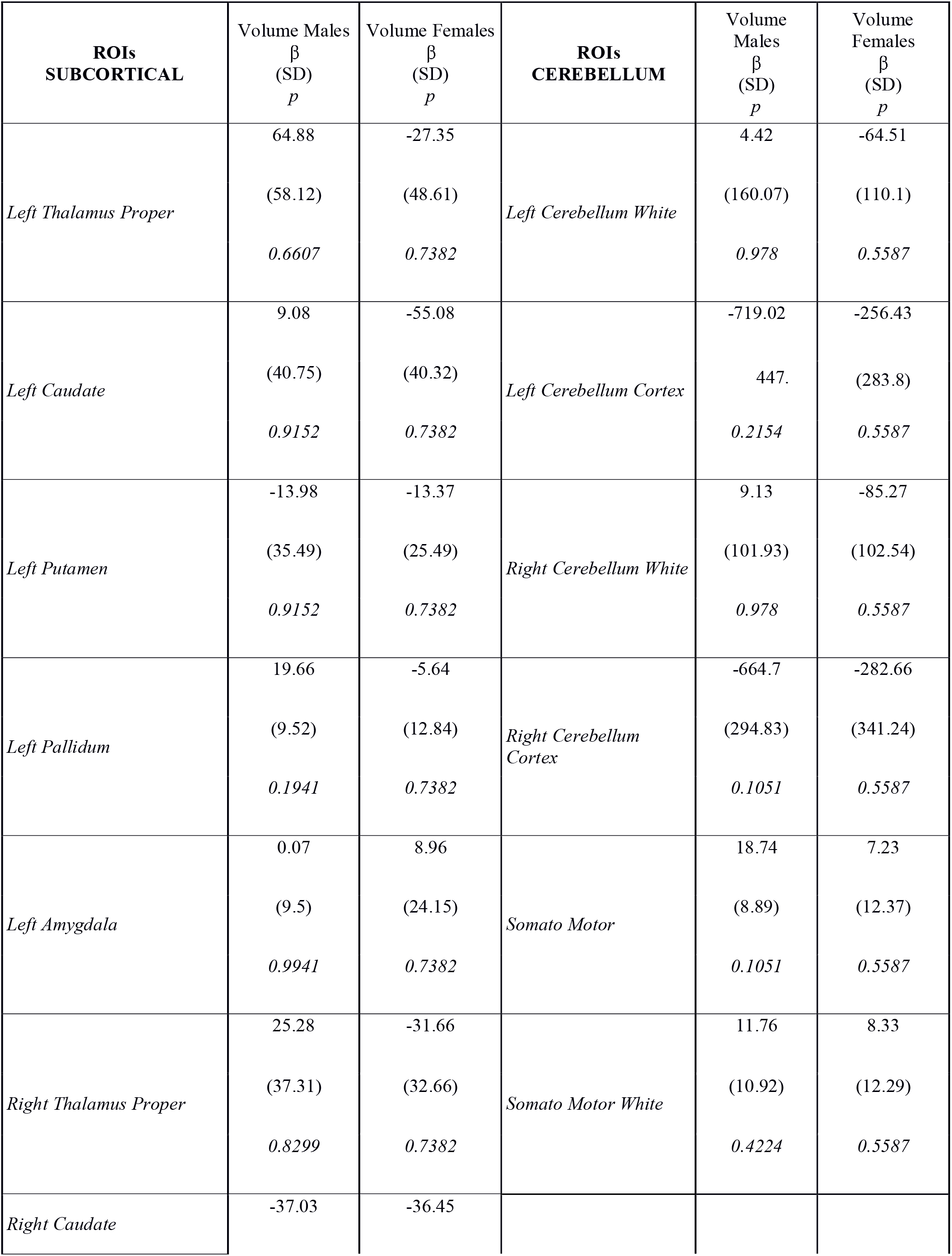
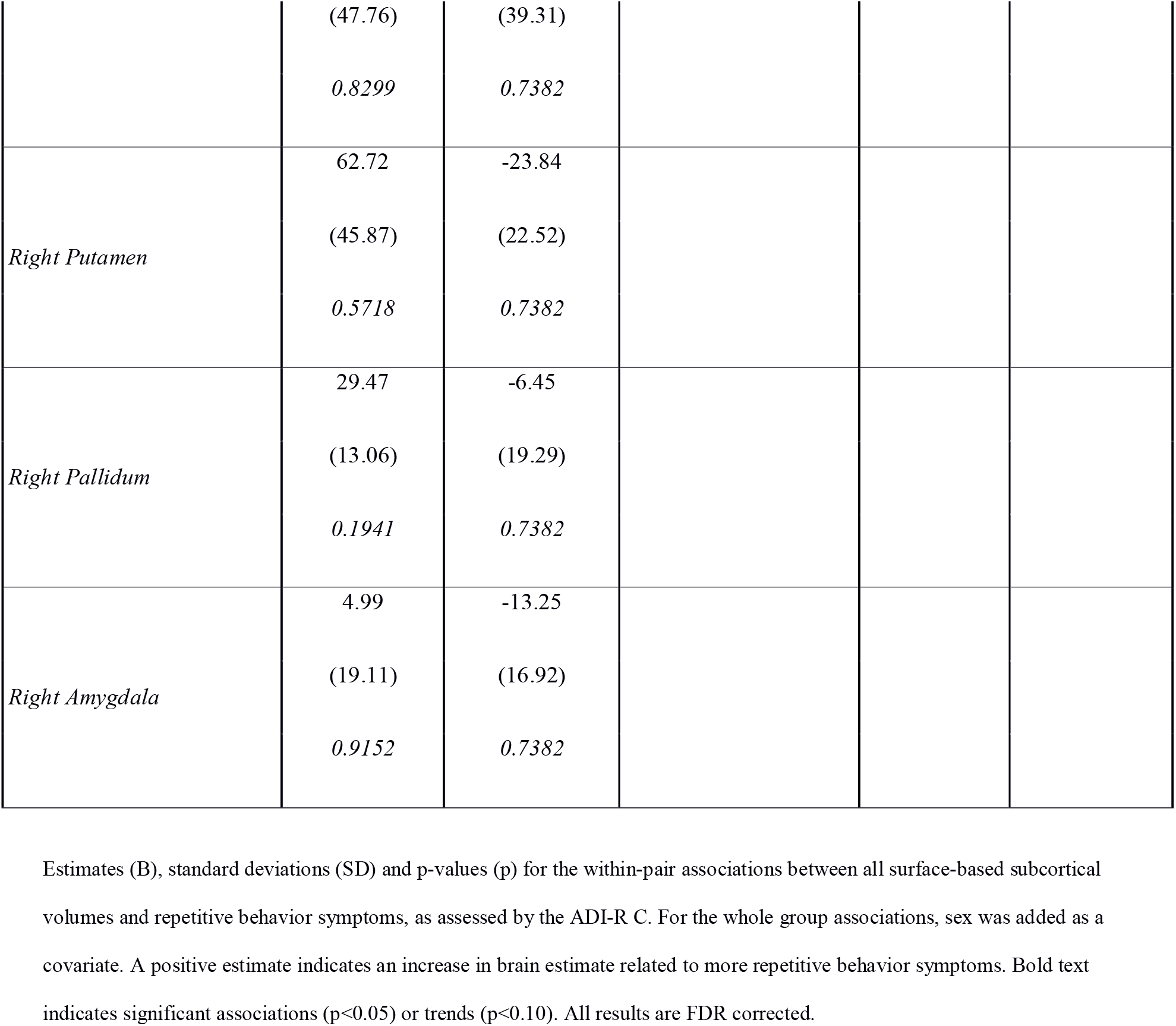
Twin model associations between subcortical volumes of subcortical regions of interest (ROIs) and RRBI symptoms.

Moreover, when controlling for non-RRBI autism symptoms and traits on the ADI-R, to test the specificity of the observed associations in females, increased thickness of the right intraparietal sulcus was still significantly associated with RRBIs on ADI-R. In addition, in females, increased thickness of the right postcentral sulcus, and increased volume of the right orbital and postcentral gyri was associated with more RRBI symptoms, whilst controlling for other autism symptoms. For males, RRBIs were associated with reduced volume of the right cerebellum cortex.

#### 2.2 Sex specific regional alterations

Further, the relationship between RRBIs on the ADI-R and brain structure differed significantly for males and females on both thickness (χ^2^=4.55, p=0.033) and surface area (χ^2^=4.02, p=0.045) of the right intraparietal sulcus, and of thickness of the right orbital gyrus (χ^2^= 4.46, p=0.035).

#### 2.3 Sex-specific results: testing robustness and specificity of the effects

In females, within-pair increases in current RRBIs, as assessed by SRS-2 RRB subscale, were associated with within-pair increases in thickness of the left intraparietal and lateral orbital sulci as well as right orbital gyrus, and increased surface area of the right supramarginal gyrus. In males, within-pair increases in current RRBIs, were associated with increased volume of the right pallidum. In addition, in males there were trends for within-pair reduction of volume and surface area of the right postcentral sulcus. However, when controlling for current social cognition impairments, these associations were no longer present (Supplementary Tables).

## Discussion

The present twin-study is the first to assess sex differences in the anatomy of brain networks associated with RRBI symptoms in autism. Significant associations were observed only within female pairs with largely varying frequencies and severities of RRBI symptoms and traits. In particular, the female twin with more RRBI symptoms had increased thickness of the right intraparietal sulcus and right orbital and lateral orbital gyri. Despite comparable within-pair differences in RRBIs, and comparable level of total autistic impairments, such associations with brains structure were not observed in males. Our results therefore suggest that largely gene-independent associations between RRBI symptoms and brain structure are mostly found in females.

Our observations correspond partly to the single previous study on sex differences in the neuroanatomy of the motor networks in ASD (Supekar and Menon 2015), where brain structure of motor areas predicted RRBIs only in girls. However, whilst Supekar and Menon report mostly primary motor regions, we observe sex specific associations with RRBIs in females in two regions involved in visuo-motor coordination, intention interpretation (intra-parietal sulcus) and executive function (lateral orbital gyri). These findings therefore correspond to the hypothesis that RRBIs are partly caused by differential sensory processing and difficulty switching attention (Shomstein and Yantis 2004, 2006). Sex specific effects are an indication of etiological differences underlying symptom domains of ASD in males and females. Indeed, interactions between sex and ASD diagnosis were observed for white matter connectivity density of the medial parietal lobe, of which the intraparietal sulcus is a part (Irimia et al. 2017). However, in that study, the sex specific effects were found only for white, not gray matter. Furthermore, abnormal intraparietal sulci (Auzias et al. 2014) and reduced gyrification of an area near the intraparietal sulcus (Pappaianni et al. 2017) were observed in male children with ASD. Functional activation studies have found differential activation in males with ASD in the intraparietal sulcus during motor learning (Travers et al. 2015) response shifting (Shafritz et al. 2008) tasks, and in the orbital gyri during motor-inhibition (Schmitz et al. 2006) and temporal delay decision making (Murphy et al. 2017), and these differences correlated in most cases with RRBI symptoms (Shafritz et al. 2008; Travers et al. 2015; Murphy et al. 2017). However, differences in functional activation are not necessarily associated with differences in underlying structure. Studies adressing functional activation during executive function tasks in females with ASD are needed in order to elucidate the relationship between structure and function in males and females with ASD. Indeed in males, RRBIs have mostly been associated with increased volume of the basal ganglia (Sears et al. 1999; Hollander et al. 2005; Haznedar et al. 2006; Rojas et al. 2006; Langen et al. 2007; Schuetze et al. 2016), thalamus (Lin et al. 2015), amygdala and precentral gyrus (Rojas et al. 2006). We do not replicate these results at our within pair level, at the exception of increased volume of the right pallidum in males when using current as opposed to lifetime presence of RRBI symptoms. Regions found in previous studies might in fact reflect rather gene dependent anatomical variation.

Other than inherent etiological differences between males and females, an explanation of our sex specific findings could be that the variance in brain structure within pairs was greater for females, therefore leading to significant associations in females but not males. Such increased brain structure differences, in combination with comparable differences in RRBI symptoms themselves, suggest greater cerebral and behavioral impairment in females for similar symptom levels. This observation might be a consequence of camouflaging. This entails that females need to have more severe RRBIs before even being noticed by their environment. Camouflaging leads to an underestimation of the true severity of autistic symptoms in females, which has recently been shown experimentally for social communication (Lai et al. 2016). In fact, females might have different types of restricted interests that might be regarded by caregivers as less atypical (Hiller et al. 2014). Thus, the true symptom levels of the females in our sample might have been higher than actually scored, which could in turn be linked to stronger or different brain anatomy alterations. Therefore, only in the most impaired females, differences in brain structure are found. Replication of our results in independent samples with assessments of explicitly high sensitivity to RRBIs in females is therefore desirable.

Related to this, an additional alternative explanation might be that the observed reductions in volume in females are related to a more general and unspecific severity of autism symptomatology, rather than to RRBIs specifically. However, re-running our analyses while regressing out highly correlated variance of other autism symptom domains and traits, thickness of both the intraparietal sulcus and orbital gyri were still associated with RRBI symptoms in females. Moreover, similar associations were observed when using RRBIs estimated with SRS-2. Compared to the SRS-2, assessing rather autistic-like traits in a short time frame (6 months), the ADI-R collects clinical RRBI symptoms, and we used scores reflecting lifetime behaviors. Therefore, our patterns of RRBIs findings on the ADI-R and SRS-2 might indicate certain alterations in anatomy of the intraparietal sulcus and orbital gyri that are clinically relevant and robust for current or past presence of symptoms. The longitudinal effects of RRBIs have been shown in children and teenagers even when using an estimate of RRBI at preschool age (Langen et al. 2014). Thus, a lifetime presence (“ever”) assessment of RRBI symptoms seems adequate to use for addressing its relationship with current brain anatomy.

Finally, in males, associations between ASD symptom domains and brain structure might be more driven by genetics, whilst in females shared and non-shared environmental factors play a more important role. Since our twin design assesses largely gene-independent associations, controlling for all factors shared between a pair of twins, the association between brain anatomy and RRBIs in the females was likely influenced by non-shared environmental factors. Future research is required to specifically assess which genetic and environmental factors contribute to neuroanatomical alterations in females with ASD, as well as if autistic females are more sensitive to these factors compared to males. Non-shared environmental factors could in this case also entail the repetitive behaviors themselves, which, when present at an early age, reinforce preexisting structural alterations (Langen et al., 2014). Further, directly assessing the influence of non-shared environmental factors would require a sample consisting of only monozygotic twins. Due to lack of power, we were unable to conduct meaningful analyses on the subsample of monozygotic twin pairs only, thus dizygotic and monozygotic twins were collapsed in the present study.

Lastly, it must be noted that, although our within-pair design compares twins of the same age, a wide age range could still have influenced the outcomes. For example, it has been proposed that basal ganglia volumes increase with age in ASD subjects due to presence of RRBIs, while they decrease in typically developing individuals (Langen et al. 2011). In addition, the precentral gyrus decreases linearly in volume in ASD subjects, but in a u-shaped form in controls (Greimel et al. 2013). Therefore, the within-pair difference related to RRBIs could be greater for older, as compared to younger subjects. Although males and females did not differ on average age in our sample, females with an ASD diagnosis were slightly older compared to diagnosed males. We can therefore not exclude the possibility that similar within-pair effects would be observed in older male subjects with ASD if they were tested. However, it is unlikely that the wide age-span by itself has confounded the results, since we tested the interaction between age and RRBIs on brain anatomy in a regular linear model, and observed significant results only in right postcentral gyrus, superior precentral sulcus and superior parietal sulcus, which were not associated with RRBIs in either sex.

In conclusion, this twin study shows that quantified features RRBI are mostly associated with brain anatomy alterations in females and that these associations are influenced by non-shared-environmental factors. The results add evidence to the hypothesis that there are etiological differences underlying ASD between males and females.

## Acknowledgments

Firstly, we are grateful for the participation, time, patience and cooperation of our twins and their parents in the RATSS project and especially during the scanning sessions. Secondly, we would like to thank all the colleagues and professionals involved in this big project for their help in collecting the data but also their advices and contributions, namely Charlotte Willfors, Kristiina Tammimies, Anna Lia Sacerdoti, Kerstin Andersson, Anna Lange Nilsson, Johanna Ingvarsson, Elin Vahlgren, Elzabieta Kostrzewa, Johan Isaksson, Martin Hammar, Christina Coco, Anna Råde, Lina Poltrago, Steve Berggren, Eric Zander, Andreas Fällman, Therese Lindström, Anna Lövgren, Torkel Carlsson, Soheil Mahdi and Lynnea Myers.

The study was funded by the Swedish Research Council, Vinnova, FORTE (previously called FORMAS), the Swedish Brain foundation (Hjärnfonden), Stockholm Brain Institute, Autism and Asperger Association Stockholm, Queen Silvia Jubilee Fund, Solstickan Foundation, PRIMA Child and Adult Psychiatry, the Pediatric Research Foundation at Astrid Lindgren Children’s Hospital, Sällskapet Barnavård, Jerring Foundation, the Swedish Order of Freemasons, Kempe-Carlgrenska Foundation, Sunnderdahls Handikappsfond, Gålöstiftelsen, the Innovative Medicines Initiative Joint Undertaking (grant agreement number 115300), which comprises financial contribution from the European Union’s Seventh Framework Programme (FP7 /2007 – 2013) and in-kind contributions from companies belonging to the European Federation of Pharmaceutical Industries and Associations (EU-AIMS).

